# Sewage Promotes *Vibrio vulnificus* Growth and Alters Gene Expression

**DOI:** 10.1101/2021.04.27.441721

**Authors:** James W. Conrad, Valerie J. Harwood

**Author notes:** Address correspondence to Valerie J. Harwood.

## Abstract

*Vibrio vulnificus* is a naturally-occurring, potentially lethal pathogen found in coastal waters, fish, and shellfish. Sewage spills in coastal waters occur when infrastructure fails due to severe storms or age, and may affect bacterial populations by altering nutrient levels. This study investigated effects of sewage on clonal and natural *V. vulnificus* populations in microcosms. Addition of 1% sewage to estuarine water caused the density of a pure culture of *V. vulnificus* CMCP6 and a natural *V. vulnificus* population to increase significantly, whether measured by qPCR or culture. Changes in the transcription of six virulence- and survival-associated genes in response to sewage were assessed using continuous culture. Exposure to sewage affected transcription of genes that may be associated with virulence. Specifically, sewage modulated the oxidative stress response by altering superoxide dismutase transcription, significantly increasing *sodB* transcription while repressing *sodA*. Sewage also repressed transcription of *nptA*, which encodes a sodium-phosphate cotransporter. Sewage had no effect on *sodC* transcription or the putative virulence-associated genes *hupA* or *wza*. The effects of environmentally relevant levels of sewage on *V. vulnificus* populations and gene transcription suggest that sewage spills that impact warm coastal waters could lead to an increased risk of *V. vulnificus* infections.

**Importance:** *Vibrio vulnificus* infections have profound impacts such as limb amputation and death for individuals with predisposing conditions. The warming climate is contributing to rising *V. vulnificus* prevalence in waters that were previously too cold to support high levels of the pathogen. Climate change is also expected to increase precipitation in many regions, which puts more pressure on wastewater infrastructure and will result in more frequent sewage spills. The finding that 1% wastewater in estuarine water leads to tenfold to 1000-fold greater *V. vulnificus* concentrations suggests that human exposure to oysters and estuarine water could have greater health impacts in the future. Further, wastewater had a significant effect on gene transcription and has the potential to affect virulence during the initial environment-to-host transition.

## Introduction

Billions of gallons of sewage are discharged into the environment and recreational waters in the U.S. annually as a result of storms, infrastructure failure, and chronic leaks from aging infrastructure (1). Sewage contains an abundance of allochthonous human pathogens which pose a direct risk to individuals who contact the water during recreation or other activities such commercial fishing, and also contaminate aquatic fisheries (1–4). Sanitary sewer overflows (SSOs) that often occur after heavy rains overwhelm local infrastructure, and may impact microbial communities if sewage enters local water bodies. Sewage and runoff contain high levels of dissolved organic carbon (DOC), nitrogen (N), phosphate (P), heavy metals, and sub-inhibitory concentrations of antibiotics which contribute to eutrophication and degraded water quality (5, 6). These nutrient pulses could further degrade local water bodies by stimulating the growth of autochthonous bacteria including human pathogens such as the leading cause of seafood borne illness fatalities, *Vibrio vulnificus* (7).

The presence of nutrients, heavy metals, and pharmaceuticals in sewage, and in other forms of wastewater, can cause disturbances in the local bacterial and phytoplankton populations when they are released to the environment. Algal blooms have been observed following heavy storms, or sewage discharge, and have been correlated with proliferation of *Vibrio* spp. resulting from increased DOC and other nutrients (8–10). Pathogenic *Vibrio* spp., (e.g. *V. cholerae, V. parahaemolyticus*, and *V. vulnificus*) can also proliferate following these events (8, 11, 12). In contrast, vibrio concentrations did not correlate with fecal indicator bacteria, which signal pollution from sewage and other sources of fecal contamination (e.g. birds (13)), in Apalachicola Bay, FL (14). One study correlated *V. parahaemolyticus* levels with the amount of wastewater treatment plant (WWTP) effluent released into Narragansett Bay, RI (15).

*V. vulnificus* is an opportunistic human pathogen that is closely related to the pathogens *V. cholerae* and *V. parahaemolyticus* (16). Humans are typically infected with *V. vulnificus* after eating contaminated oysters, which can result in septicemia and up to a ~50% mortality (17). Exposure of wounds to estuarine water or animals (e.g. shellfish or fish) can result in cutaneous infections and necrotizing fasciitis, which may necessitate limb amputation (17). Naturally-occurring *V. vulnificus* populations consist of three major biotypes; biotype one causes the majority of human infections (18, 19). Within biotype one, *V. vulnificus* is grouped into environmentally-associated (16S rRNA A or *vcgE*) and clinically-associated (16S rRNA B or *vcgC*) genotypes. The 16S rRNA A/B and *vcgC/E* typing methods are both used frequently and have a high degree of concordance (20–23). The clinically associated-genotype 16S rRNA B genotype is more frequently isolated from human infections and is correlated with more severe disease outcomes compared to the environmentally-associated 16S rRNA A genotype (20, 23, 24). Differential expression of genes by each genotype may contribute to the observed genotype bias in clinical specimens. The sodium phosphate cotransporter *nptA* is differentially expressed by *V. vulnificus* genotypes (25) and may support growth under changing phosphate concentrations as observed in *Staphylococcus aureus* (26).

Expression of virulence genes in bacteria has been shown to respond to environmental conditions including temperature (27–29), salinity (25, 30), carbon sources (31–33), nutrients (25), heavy metals (34), and antibiotics (33, 35, 36). Sewage represents a source of numerous organic carbon molecules (37) inorganic nutrients, and metals (38). Iron is found in high concentrations in wastewater and can be a limiting nutrient in seawater for algae (39, 40), but also is potentially toxic, inducing oxidative stress in bacteria (41, 42). *hupA* expression in *V. vulnificus* is important for iron acquisition during infections (43). Antioxidant-related changes in gene expression (e.g. *sodA-C*) can promote survival and virulence under acid stress and phagocyte engulfment in *V. vulnificus, V. alginolyticus*, and *Salmonella enterica* (42, 44–46). Changing nutrient levels, resulting from sewage, can affect the expression of genes related to nutrient acquisition and contribute to virulence potential. Similarly, expression of a capsule (e.g. *wza*) increases survival of *V. vulnificus* in the presence of serum (47–49) and is affected by environmental conditions (e.g. temperature and oxygen availability) (50, 51).

Sewage could directly influence the probability of human infection by *V. vulnificus* if it stimulated growth of the pathogen. On the other hand, sewage could indirectly increase pathogen infectiousness by altering the expression of genes related to virulence and the environment-to-host transition through multiple mechanisms. This study’s purpose was to investigate the effects of sewage on *V. vulnificus* growth and gene transcription using both laboratory cultures and natural populations of bacteria present in estuarine water in Tampa Bay, FL. The objectives were to 1) determine if sewage can serve as a nutrient source for autochthonous *V. vulnificus* populations; and 2) determine if sewage alters the transcription of virulence- and survival-associated genes.

## Methods

### Strains and culture conditions

*Vibrio vulnificus* strain CMCP6 was maintained on Luria-Bertani agar (Difco). *V. vulnificus* CMCP6 broth cultures prepared for inocula in microcosm and gene transcription experiments were incubated for 20-24 h in Luria-Bertani (LB) broth at room temperature (22°C).

### Sample collection and processing

Sewage influent was collected from Falkenburg Advanced Wastewater Treatment Plant, Tampa, FL, transported on ice, and held for a maximum of two hours before being frozen at −20°C. Sewage was held in the freezer for a maximum of one month prior to thawing and filter sterilization with a Rexeed 25-S hollow-fiber filter (Asahi Kasei). Estuarine water was collected from Ben T. Davis Beach (BTD) Tampa, FL 27°58’12.9’’N, 82°34’42.9’’W (pH 7.9, salinity 16-22 ‰) and Hudson Beach, Hudson FL 28°21’46.3”N 82°42’33.6”W (pH 7.8, salinity 20 ‰) and used to construct microcosms, or sterilized by hollow fiber filtration as above and frozen, within 4 h of collection.

### Assessing the effects of sewage on growth of *V. vulnificus* CMCP6

The ability of sewage to serve as a nutrient source was assessed by incubating *V. vulnificus* CMCP6 in microcosms with and without sewage. *V. vulnificus* concentrations were measured by qPCR of the *vvhA* gene (Table 1) (52). All microcosms were prepared in triplicate. *V. vulnificus* CMCP6 inoculum was grown at room temperature for ~22 h in LB broth and diluted to ~10^3^ CFU/mL in phosphate buffered saline (pH 7.4) (53). A 100 μL aliquot of diluted culture was added to each 20 mL microcosm to reach a starting concentration of ~10^1^ CFU/mL and incubated at 37°C with shaking at 150 rpm for ~22 h.

**Table 1.**
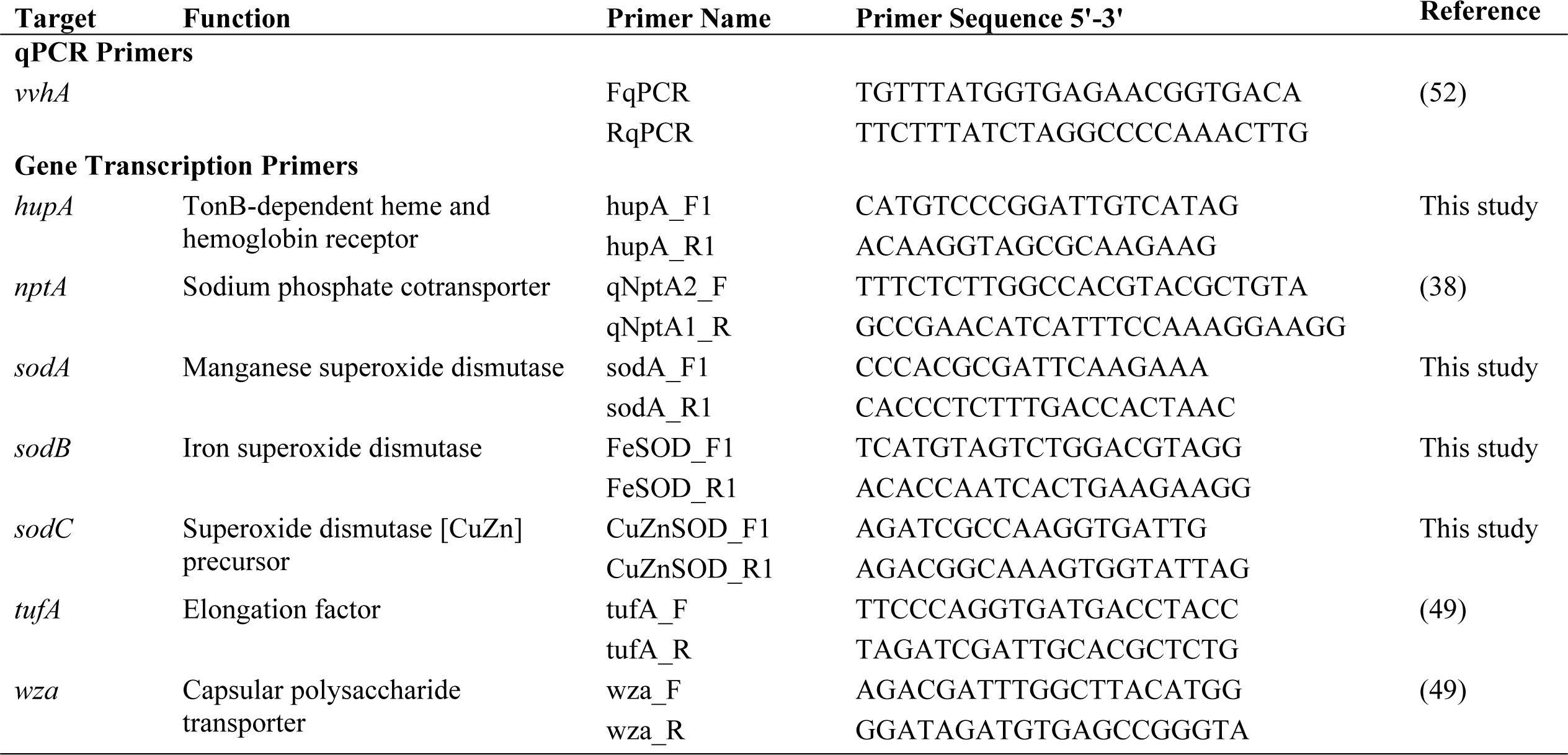
qPCR and RT-qPCR primers used in this study.

Effects of the macronutrients nitrogen, phosphorous, and organic carbon in sewage on *V. vulnificus* CMCP6 growth were investigated by preparing a defined modified M9 (MM9) medium lacking these macronutrients. MM9 was amended with sterile sewage to serve as the sole source of the missing nutrients to determine their effects on culture density. Control (nutrient-replete) microcosms contained 20 ml of MM9 media consisting of 50 mM tris HCl (pH 7.5), 10 mM NH_4_Cl, 0.1 mM CaCl_2_, 1 mM MgSO_4_, 1 mM KH_2_PO_4_, 0.1 mM ferric citrate (C_6_H_5_FeO_7_), 10 ‰ NaCl, and 11.1 mM (2 g/L) glucose. A medium depleted of nitrogen, phosphorous and carbon was prepared by omitting NH_4_Cl, KH_2_PO_4_, and glucose from MM9. Estuarine water from Hudson Beach, FL (pH 7.8, salinity 20 ‰) was sterilized using hollow-fiber filtration (Rexeed 25-S) for microcosms made with environmental water. Sewage-amended treatments received 1% (vol/vol) sterile sewage influent. An undiluted sterile sewage treatment was amended with NaCl to a salinity of 10 ‰. Twenty milliliters of culture, 1 mL for turbid cultures, were filtered through a 0.45 μm nitrocellulose filter to concentrate bacteria. Membrane filters were stored at −80°C until DNA could be extracted using a DNeasy Power Water kit (Qiagen) and bacteria were quantified using qPCR of the *vvhA* gene.

### Assessing the effect of sewage on the growth of autochthonous *V. vulnificus*

The effects of nutrient amendment on culturable concentrations of autochthonous *V. vulnificus* populations in estuarine water was assessed in batch cultures. Microcosms (500 mL) were constructed in triplicate using estuarine water from BTD (pH 7.9, salinity 22 ‰). We used a control treatment (natural estuarine water only), a low-level glucose amendment (3.0 mg/L glucose)(54), and a sewage amendment (1% filter-sterilized sewage influent). Microcosms were incubated at 30°C with shaking at 140 rpm for 20-24 h. Culturable *V. vulnificus* were enumerated using membrane filtration by filtering 1 mL of serially diluted culture through 0.45 μm nitrocellulose membrane filters and plating on modified cellobiose-polymyxin b-colistin agar (mCPC) (55). Plates were incubated at 37°C for 22-24 h and then counted.

### Effects of sewage on virulence- and survival-associated genes

Changes in transcription of six virulence- and survival-associated genes (*hupA, nptA, sodA-C, tufA*, and *wza*) by *V. vulnificus* CMCP6 in response to sewage were assessed using an Infors-HT II bioreactor in a chemostat (continuous flow) configuration. Genes were selected based on their potential dual roles in survival in the environment and the human host, or known importance for virulence expression. Defined minimal medium containing 23.3 mM Na_2_HPO_4_, 11 mM KH_2_PO_4_, 9.35 mM NH_4_Cl, 85.6 mM NaCl, 1 mM MgSO_4_, and 2.25 mM glucose (0.405 g/L) with pH adjusted to 7.5 was used as a growth medium. The 1 L culture vessel and 4 L reservoir were filled with medium and the bioreactor was set to pH 7.5, temperature 37°C, dissolved oxygen >70%, 150 rpm stir rate, and a flow rate of 3 L/d. The *V. vulnificus* CMCP6 inoculum was grown at room temperature for ~22 h in LB and 1 mL of culture was added to the bioreactor to reach a starting concentration of 10^6^ CFU/mL After inoculation, the bioreactor was run continuously for 48 h prior to sampling under control (no sewage added) conditions. Sampling under control conditions occurred thrice over the course of 4 h. After sampling under control conditions, the nutrient reservoir was replaced with minimal medium amended with 1% (vol/vol) sterile sewage and allowed to run for another 48 h. After 48 h, the sewage treatment was sampled thrice over the course of 4 h.

Immediately after each sample collection RNA was extracted using a Quick-RNA Miniprep Kit (Zymo) followed by a dnase treatment using a TURBO DNA-free Kit (Invitrogen). Briefly, RNA was diluted to 10 ng/μL and used for reverse transcriptase qPCR (RT-qPCR) of the following genes: *hupA, nptA, sodA-C, tufA*, and *wza* (Table 1). Thermo Scientific™ Verso™ 1-Step RT-qPCR Kits with low ROX (Thermo Scientific) was used for one step reverse transcription in an ABI 7500 qPCR thermocycler. Twenty microliter qPCR reactions consisted of 1x Verso master mix, 1 μL enhancer per reaction, 0.2 μL Verso Enzyme per reaction, 0.15 μM of each primer (Table 1), 2 μL of template RNA (10 ng/ μL) per reaction, and nuclease free water. Cycling conditions were as follows: 1 cycle of 50°C for 15 min followed by 1 cycle of 95°C for 15 min followed by 40 cycles of 95°C for 15 s and 60°C for 30 s. Dnase treatment was verified using a no enzyme control (reactions lacking Verso Enzyme). Fold gene transcription was calculated using the 2^−ΔΔC^_T_ method, which normalizes transcription to a reference gene, (56) with *tufA* serving as the reference gene (57).

### Statistical analyses

Statistical analyses on culturable bacterial concentrations, qPCR, and RT-qPCR data were performed in R v3.6.3 and Graphpad Prism v8. ANOVA followed by Tukey’s honest significance tests was performed using Graphpad and the package multcomp in R.

## Results

### Sewage promotes *V. vulnificus* CMCP6 growth

We sought to determine if sewage could support the growth of *V. vulnificus* CMCP6 in a minimal medium and in sterile estuarine water. Culture density in microcosms containing nutrient replete MM9 (4.67 × 10^9^ GC/100 mL) was not significantly different from MM9 with 1% added sewage (5.47 × 10^9^ GC/100 mL) or from cultures grown in undiluted sewage (5.33 × 10^9^ GC/100 mL) (Fig. 1). *V. vulnificus* CMCP6 in nutrient depleted MM9 (lacking a nitrogen, phosphorus and carbon source) amended with 1% sewage (NPC lim + 1% Sew) reached a density of 6.81 × 10^7^ GC/100 mL, while *V. vulnificus* concentrations in NPC-depleted medium without sewage were below the limit of detection (< 10 GC/mL) (data not shown). The addition of 1% sterile sewage to sterile estuarine water caused a significant 1.16 log_10_ GC/100 mL increase in *V. vulnificus* density to 4.21 × 10^7^ GC/100 mL compared to the sterile estuarine water (2.88 × 10^6^ GC/100 mL) (Fig. 1). *V. vulnificus* levels in nutrient-depleted MM9 amended with sewage were not significantly different than those in sterile estuarine water amended with sewage.

**Figure 1.**
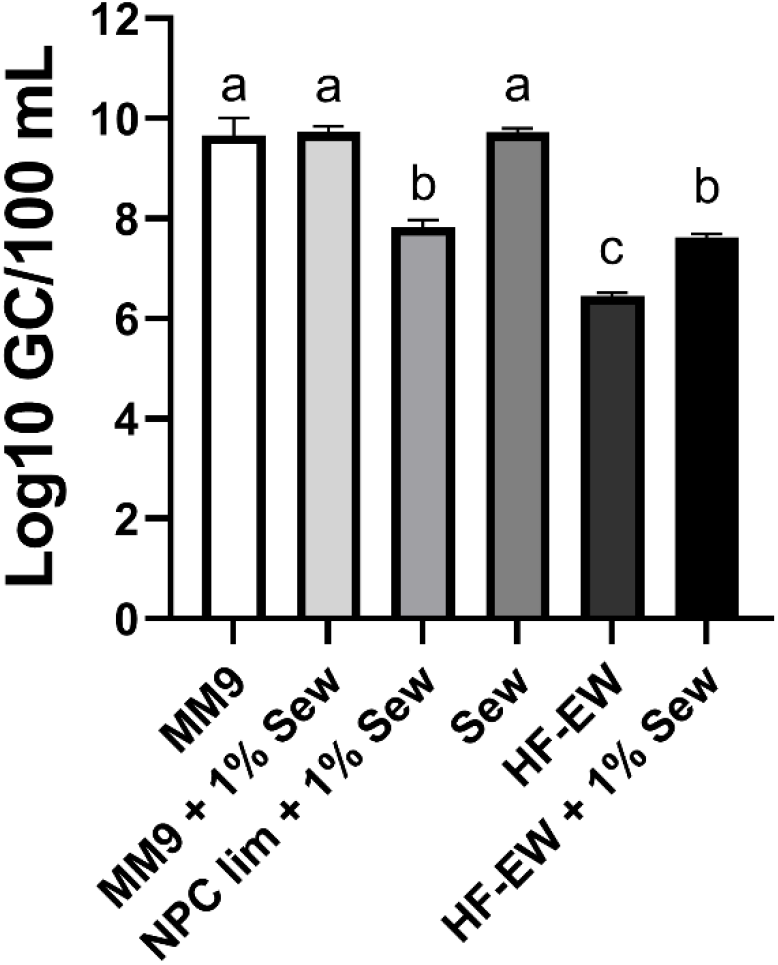
Effects of sewage on the density of *V. vulnificus* CMCP6 measured by qPCR of *vvhA*. *V. vulnificus* CMCP6 was grown in the following media with or without 1% sterile sewage added: nutrient replete minimal media (MM9), MM9 without added nitrogen, phosphorous, and carbon (NPC lim), and sterilized estuarine water (HF-EW). It was also grown in undiluted sterile sewage (Sew). Treatments listed with “+ 1% Sew” received a 1% (vol/vol) sterile sewage amendment to growth medium. *V. vulnificus* density in the NPC limited media without sewage was below the limit of detection (not shown). Error bars represent the standard deviation of the mean between replicates and letter codes indicate a significant difference between treatments when letters are not shared (p ≤ 0.05).

### Sewage supports the growth of natural *V. vulnificus* populations

We explored the potential for sewage to increase the density of a natural population of *V. vulnificus* by incubating autochthonous populations in natural estuarine water + 1% sterile sewage for 24 h (Fig. 2). The effect of 3.0 mg/L (16.7 μM) glucose, used to simulate organic carbon resulting from primary production, on the growth of *V. vulnificus* was also examined. The autochthonous *V. vulnificus* population grew to a significantly greater density as measured by culture in the sewage-amended microcosms in 24 h; i.e. 2.17 × 10^6^ CFU/100 mL in the sewage treatment compared to 8.49 × 10^2^ CFU/100 mL in the un-amended estuarine water (Fig. 2). Added glucose caused no significant difference in culturable *V. vulnificus* concentrations compared to the un-amended estuarine water (Fig. 2).

**Figure 2.**
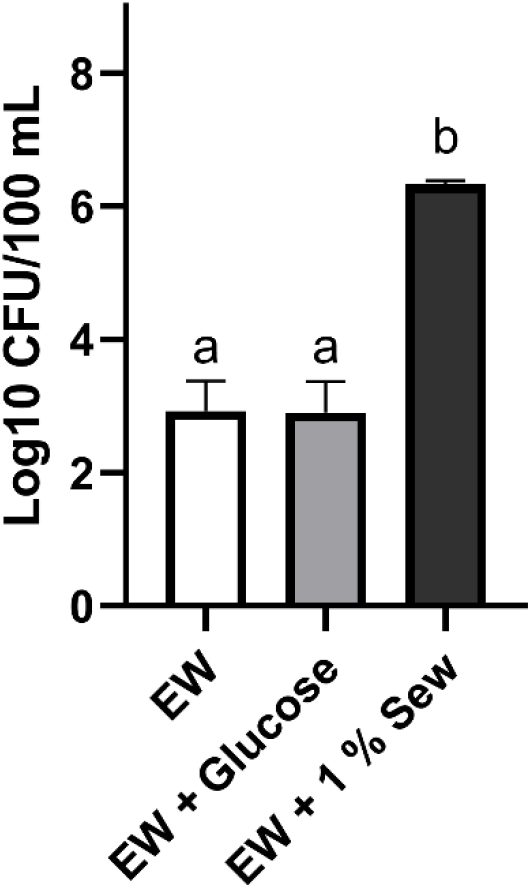
Density of an autochthonous *V. vulnificus* population measured by culture after 24 hours of growth in natural estuarine water (EW) from Ben T. Davis Beach, FL. Treatments were unamended EW, EW amended with 3.0 mg/L glucose (EW + Glucose), and EW amended with 1% sterile sewage (EW + 1% Sew). Error bars represent the standard deviation of the mean between replicates and letter codes indicate a significant difference between treatments when letters are not shared (p ≤ 0.05).

### Effects of sewage on gene transcription

The possibility that compounds in sewage could affect the transcription of virulence- and survival-associated genes was tested using *V. vulnificus* CMCP6. *V. vulnificus* CMCP6 was maintained as an actively growing culture using a bioreactor in a chemostat configuration with nutrient replete medium. A stable continuous culture was established and sampled before being exposed to 1% sewage to determine changes in the transcription of virulence- and survival-associated genes (*sodA-C*, *hupA, nptA*, and *wza*). Sewage exposure significantly increased Fe SOD (*sodB*) transcription 2.7-fold over the control (Fig. 3). Conversely, transcription of *sodA*, which encodes Mn SOD, significantly decreased 5.4 fold upon exposure to sewage. *nptA* transcription was a significant 2.1-fold lower in the sewage treatment compared to the control. Changes in transcription of the remaining genes (*sodC* encoding the CuZn SOD, *hupA*, and *wza*) were not significant. While not significant at α=0.05, sewage appeared to repress *hupA* (p = 0.08) transcription.

**Figure 3.**
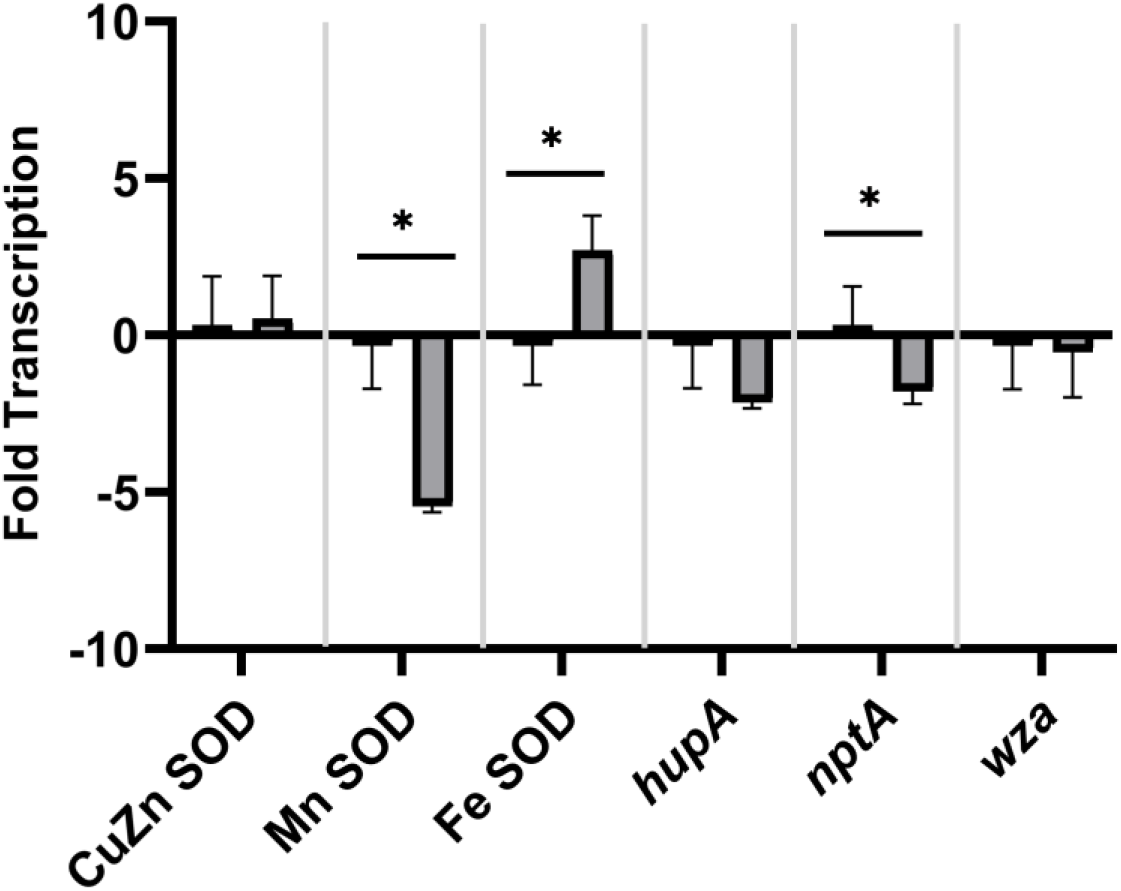
Changes in fold-transcription of virulence- and survival-associated genes in response to amendment with 1% sewage was assessed by RT-qPCR: *sodC* (CuZn superoxide dismutase (SOD)), *sodA* (Mn SOD), *sodB* (Fe SOD), *hupA*, *nptA* and *wza*. Cultures were grown using a bioreactor in unamended minimal medium (control, □ on left) or in minimal medium + 1% sterile sewage (sewage, ■ on right). Error bars represent the standard deviation of the mean between replicates and asterisks represent a significant difference in the mean between treatments (with or without sewage) (p ≤ 0.05).

## Discussion

Contamination of surface waters by sewage and storm water is known to endanger human health by increasing the probability of human exposure to allochthonous pathogens, and also to degrade water quality through nutrient loading (58–60); however, the possibility that sewage promotes increased levels of autochthonous aquatic pathogens by providing nutrients has been infrequently addressed. Sewage is often accidentally discharged into the environment during heavy rains where storm drains and sewer systems are connected, or where leakage from aged septic and sewer systems occurs, resulting in millions of gallons of sewage contaminating the environment annually (61, 62). Demonstrated increases in *Vibrio* spp. concentrations following storm events have been attributed to reduced salinity and mixing of shallow and deep waters (8, 12, 63). However, the effects of sewage on autochthonous, pathogenic *Vibrio* spp. (e.g. *V. cholerae*, *V. parahaemolyticus*, and *V. vulnificus*) are largely unexplored and may represent a threat to human health, as higher concentrations of pathogenic *Vibrio* spp. significantly increase the risk of infections (64).

This study demonstrated that environmentally relevant sewage levels can significantly increase *V. vulnificus* density. The concentration of 1% sewage used here was selected as it represents a reasonable level of contamination following a recent, local sewage spill or chronic contamination. We base this assessment from a review of human associated *Bacteroides* genetic marker (HF183) which is commonly used to identify sewage contamination of surface waters (58, 65). HF183 levels of 6.31 × 10^5^ - 6.15 × 10^6^ GC/100 mL have been measured in sewage diluted 100-fold (1%) (66–68) which is within the range of 1.80 × 10^3^ - 6.30 × 10^7^ GC/100 mL observed in moderately to severely impacted surface waters (13, 68–71). A low level of organic carbon (3.0 mg/L) was tested to simulate the level of organic carbon from primary production in an estuary (mean 3.07 mg/L) (54) but did not affect the observed population density in this study.

We demonstrated that sewage promotes proliferation of both pure cultures of *V. vulnificus* CMCP6 and natural *V. vulnificus* populations. Growth of *V. vulnificus* CMCP6 in sterile estuarine water without sewage resulted in a population density of ~10^6^ GC/100 ml, which is at the upper level of previous reports from Gulf of Mexico coastal waters (72, 73). The addition of 1% sewage in this study increased CMCP6 density by an order of magnitude, bringing it above the range observed in the aforementioned reports. Likewise, levels of natural *V. vulnificus* populations measured by culture in this study (~10^3^ CFU/100 ml) were similar to previously observed levels (72), but, with the addition of sewage, increased over three orders of magnitude to levels rarely reported in environmental waters. The increase in natural populations associated with sewage amendment was corroborated in a continuous culture experiment where *V. vulnificus* was measured by qPCR of *vvhA* (74). *V. vulnificus* concentrations increased over one hundred fold in 24 h following sewage addition, from ~10^5^ GC/100 ml to ~10^7^ GC/100 ml. While one would expect qPCR measurements to be higher than culture measurements, the magnitude difference in the effect of sewage among the different experiments was unexpected. It is possible that measurements of density of the autochthonous population by culture underestimated the initial quantity of *V. vulnificus* due to the presence of viable but nonculturable *V. vulnificus*, but the addition of sewage not only promoted proliferation, but also shifted a greater proportion of the cells to culturability (52).

Based on the growth-promoting effects of sewage, and presence of bioactive compounds, we hypothesized that sewage would induce the transcription of several virulence- and survival-associated genes which may facilitate the environment-to-host transition. Sewage represents a rich source of iron with concentrations ranging from 1.9-17.3 mg/L to >70000 mg/kg in sludge (38, 75). Elevated *sodB* (Fe SOD) transcription and *sodA* (Mn SOD) repression observed here is consistent with *fur*-mediated gene regulation in the presence of iron observed in *V. vulnificus* and *Escherichia coli* (44, 76). Fe SOD expression has been shown to be more important for virulence expression in mice than either *sodC* (CuZn SOD) or *sodA* (42). Expression of Fe SOD was also demonstrated to be an important virulence factor in fish infections in the opportunistic human pathogen *Vibrio alginolyticus* (46). Elevated transcription of *sodB* may facilitate the environment-to-host transition and could be an important virulence factor in human infections. While not observed here, *hupA* transcription increases upon exposure to human serum, allowing for iron acquisition (77). Free iron provided by sewage may have repressed transcription of *hupA* (29)*. nptA* transcription was repressed in response to sewage. Phosphorus concentrations in sewage are approximately three orders of magnitude higher (3 mg/L or 31.6 μM (78)) than those in estuarine water in Florida Bay (0.02-0.04 μM (79)), which could explain the effect on phosphate transporter transcription. However, it was reported that phosphate concentration does not affect *nptA* (25), and it is possible that other or multiple constituents of sewage contributed to the observed effect. While the function of *nptA* in *V. vulnificus* pathogenesis is not well understood, its expression under varying environmental conditions (25) may support the transition to a human host, as proposed for *nptA* expression in *Staphylococcus aureus* (26), by enabling rapid phosphate uptake in the new environment.

This study has shown that sewage represents a threat to human health beyond direct deposition of allochthonous pathogens. Sewage can alter the autochthonous *V. vulnificus* population in multiple ways by stimulating growth and increasing the transcription of multiple virulence associated genes. The response of *V. vulnificus* and other pathogenic *Vibrio* species to sewage may also enable better modeling of human health risks. Studies comparing opportunistic pathogens to obligate pathogens will be important to understand the broader impacts of sewage on waterborne pathogens and risk to human health.

## Acknowledgements

We thank Dr. Anita Wright for providing us with a *V. vulnificus* CMCP6 culture, funding from the Porter Family Foundation (USF), and the Aylesworth Scholarship (USF).

